# Connectivity and Contraction in Cytoskeletal Networks

**DOI:** 10.1101/2025.06.12.659219

**Authors:** Michael J. Norman, Annamarie Leske, Julio M Belmonte

## Abstract

The cytoskeleton”s ability to contract and propagate forces is the fundamental mechanism behind cell morphology, division and migration. This can only occur if the network is sufficiently connected, yet a rigorous description of the connectivity requirements has never been provided. In this work we focused on the polarity-sorting contraction mechanism and showed that connectivity is not determined by the spatial distribution of filaments alone, but by the interconnectivity between the dual network of filaments and motors. We developed a method to quantify filament-motor connectivity as a function of motor length, filament length distributions, and the densities of each component. Using this framework, we derived a general theory that predicts when a network is sufficiently connected to allow global or local contraction. We validated our predictions with computer simulations and introduced a novel metric to distinguish between these outcomes. Our findings show that the conditions for local and global contraction in the presence of fiber dynamics correspond, respectively, to the pulsatile and steady-state contraction behaviors observed in vivo. All results are independent of filament rigidity, making our conclusions applicable to both actin and microtubule networks. Lastly, we discuss how those outcomes are affected by the introduction of crosslinking proteins, which - despite not actively generating forces of their own - can promote global contractility at small concentrations even in networks made of short and/or rigid filaments.

## INTRODUCTION

The development of multicellular organisms involves changes of tissue structure, where groups of cells differentiate, change shape and move to form new tissues or organs. All these processes depend on the generation of intracellular forces and on the transmission of those forces across the cells and tissues. One of the main sources of those forces is the actin cytoskeleton, a system of proteins, molecular motors and fibers linked to the plasma membrane (**Fig 1A**). Disruption of any component of the actin cytoskeleton can affect many cellular processes such as migration (Yamada 19, Blaser 06), division (Leite 19), and tissue-scale processes such as gastrulation, convergent-extension and cell sorting (Clarke 21, Kim 11, Martin 09, Maître 12). On a microscopic level, the method through which a single molecular motor consumes energy and generates force on a single fiber is relatively well understood. On a macroscopic level, however, the generation of net contractile rearrangement of fibers depends on external and geometrical factors that modulate its function and ensures that the forces act across the whole network. Without a sufficient density of cytoskeletal components, cellular structures like cytokinetic rings, ring channels, cell cortices, and supra-cellular actin networks –as those found during multicellular development – cannot work.

**Figure 1.**
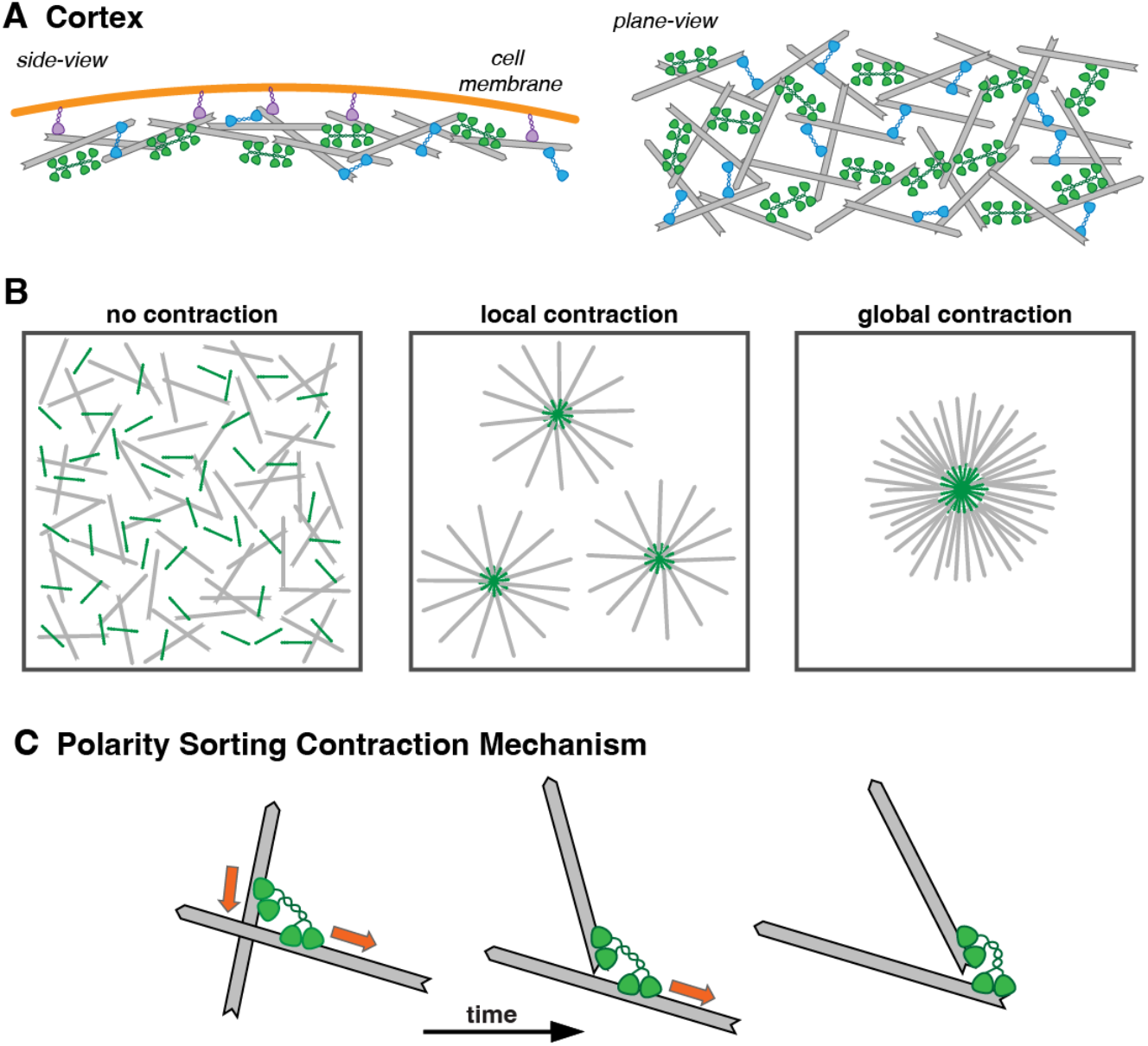
Actin cytoskeleton network composition and contraction. A) Actin network in cell cortex seen in cross-section and in plane view. Basic elements of actin networks include fibers (gray), myosin minifilaments (green), crosslinkers (blue) and membrane-to-cortex anchors (purple). B) Three possible contraction outcomes. C) Polarity sorting occurs when multivalent motors can dwell at one fiber end while processively walking along another, bringing both fibers” end together.

A large body of experimental research has investigated how cytoskeletal network connectivity governs contraction, force transmission, and structural organization. Many studies use in vitro reconstituted actomyosin systems with varying combinations of actin, myosin, and crosslinkers. For example, Bendix *et al*. (Bendix 08) and Alvarado *et al*. (Alvarado 13) found that global contraction only occurs within specific ranges of myosin and crosslinker concentrations, with too few or too many of either preventing contractility. Köhler *et al*. (Köhler 12) demonstrated that crosslinker type is critical: polar crosslinkers like fascin enable robust and repeated contraction with low concentrations, while apolar crosslinkers like cortexillin require much higher levels to achieve similar effects. Similarly, Ennomani *et al*. (Ennomani 16) showed that the organization of actin filaments influences connectivity—disorganized rings fail to contract unless crosslinkers are added.

Other studies highlight the role of motors as structural linkers. Linsmeier *et al*. (Linsmeier 16) showed that actomyosin networks can contract without crosslinkers, indicating that motors alone can provide sufficient connectivity. Thoresen *et al*. (Thoresen 13) further demonstrated that different myosin variants, including non-muscle forms, can regulate actin bundling and contraction depending on motor properties like length and stall force. In networks with fiber turnover, Krishna *et al*. (Krishna 24) found that both motor and fiber densities must be high enough to maintain continuous contraction, with patterns varying based on those concentrations.

Many previous researchers have sought to understand the relationship between network connectivity and global contractile behavior at a theorical level. As early as in 1995, Forgacs (Forgacs 95) discussed the role of percolation in biological contexts and cites the cytoskeleton network as an example of a system that needs to be sufficiently connected in order to propagate information within and between cells. In Alvarado *et al*. (Alvarado 17), the authors review many approaches to investigating the role connectivity plays in network contraction and highlight how approaches that aim to connect regular grid percolation theory with gel experiments may be useful for understanding certain macroscopic material properties. They discuss how connectivity needs to be viewed in terms of both motor and crosslinker densities and how properties like protein length and motor speed will affect connectivity and contraction.

Computational studies have employed a range of models—from lattice-based to agent-based simulations—to investigate how cytoskeletal network connectivity governs force transmission and contractility. Many of these models aim to reproduce the threshold behavior observed experimentally, where contraction only occurs if motor activity or connectivity exceeds a critical level. Wang *et al*. (Wang 12) and Frolov *et al*. (Frolov 24) used node-based and agent-based models, respectively, to show how myosin-generated forces can drive local and global contraction, with insufficient motor or connectivity levels resulting in stalled or fragmented networks. Bueno *et al*. (Bueno 22) developed a mean-field theory to predict percolation thresholds based on motor and crosslinker densities, offering a high-level framework that aligns with observed transitions between non-contractile and contractile states,

Several studies used structured grids or lattices to simplify analysis. Sheinman *et al*. (Sheinman 12) and Kumar *et al*. (Kumar 23) placed fibers on cubic and hexagonal lattices, respectively, examining how the percentage of filled edges and motor strength influence elasticity and stress propagation. Wang *et al*. (Wang 24) introduced anisotropic connectivity in a triangular grid, showing direction-dependent percolation thresholds that may reflect cytoskeletal alignment. Similarly, Ueda *et al*. (Ueda 23) used a square lattice to model stress fiber formation and found that fiber length distributions significantly influence percolation and force propagation.

Other models emphasize the role of component interactions. Zeng *et al*. (Zeng 11) used a 3D spring-based model to represent cytoskeletal elements, focusing on stress transmission from membrane to nucleus. Linsmeier *et al*. (Linsmeier 16) developed both gel-based and agent-based models to study contractility without crosslinkers, ultimately identifying density and motor activity as the key drivers. Ennomani *et al*. (Ennomani 16) combined experimental work with simulations to show how crosslinkers enable local myosin forces to produce global contraction, though they did not account for motors” own linking function in their connectivity definitions.

All these works underscore the complexity of translating percolation concepts into biologically realistic models. Although many authors employed the term *connectivity* in explaining their results, the term is never clearly defined, and its use differs greatly between groups. Work like Bendix, Köhler and Ennomani focus on the connectivity brought by the activity of crosslinkers only, while overlooking the fact the motors also contribute to connection alongside their walking activity, as shown by the work of Linsmeier *et al*. (Others, as in Bueno *et al*. (consider both actin and motor concentrations but ignore the role that fiber and motor lengths may have on connecting a network. While these models offer valuable theoretical insights, most used idealized structures such as regular lattices or randomly connected nodes, so their predictions about thresholds and force propagation must be interpreted cautiously as it is not immediately clear how to translate these abstract graph or lattice structures into the complex, disordered architecture of real cytoskeletal networks.

In this work, we present a theory that attempts to keep in as much about the local physical activity of the components necessary for cytoskeletal function while being general enough to account for a wide variety of outcomes. To predict global network behavior, we use the local interaction between fibers and motors as a starting point. The simplest and most common method through which motors can reorganize is known as polarity sorting (**Fig 1C**), a process by which multivalent motors can dwell at the end of a filament while still processivity walking along another, thus bringing each filament ends together (Kruse 00, Nedelec 01). Many microtubule motors exhibit this filament end-dwelling property (Akhmanova 05), which has shown to lead to network contraction into asters (Surrey 01, Foster 15, Torisawa 16). Actin networks have also been reported to contract into asters (Verkhovsky 97, Backouche 06, Soares e Silva 11, Luo 13, Köster 16, Stam 17), and in 2018 the end-dwelling property of myosin motors have been shown experimentally (Wollrab 18). When the network conditions are right, the network will contract into a single aster where the barbed ends of a large number of fibers are all pulled towards the same location and adjacent fibers will be closely aligned. However, at lower connectivity, the system may contract only locally, forming multiple asters, or not at all (**Fig 1B**).

Working under this contraction mechanism, we derived a theory that correctly predicts when a network will contract locally, globally or not at all. By quantifying the connectivity formed by direct geometric intersections between individual fibers and motors we show that cytoskeleton connectivity is a function of fiber and motor densities and their associated length distributions. Using local mechanical models together with a new contraction metric capable of distinguishing between local vs global contraction, we predict under what conditions contraction is impossible, possible but unreliable or guaranteed to contract the entire system into one single aster. Predictions are tested using physics-based simulations generated by the simulation software *CytoSim* where we observe how the size and spatial distribution of such networks change as a function of the number of fiber-motor crossings per motor and per fiber. We then discuss how those outcomes are affected by the introduction of crosslinking proteins, which, despite not actively generating forces of their own, can promote global contractility at small doses even for networks made of short and/or rigid filaments. We explore the effect of spatial boundary conditions on contraction behavior. Finally, we show that our results can be applied to systems with fiber turnover, which are of particular importance to the early development of organisms.

### THEORY

For local actin–motor forces to result in network-wide contraction, the cytoskeletal components must form a connected structure. For the polarity sorting contraction mechanism, connectivity can be though as the number of fiber-motor crossings present in the network. In order for this mechanism to work every motor must connect to at least 2 fibers, while every fiber must connect to at least 1 motor (**Fig 2A**). This minimal configuration ensures that the fundamental contraction mechanism is functional (**Fig 1C**) but does not guarantee global contraction as the contractile units are not necessarily connected to each other. To ensure a global connectivity and contraction, we hypothesize that every fiber must be connected to at least 2 motors (**Fig 2A”**). When these conditions are met throughout the network, any given fiber that is being pulled by a motor will also pull any other motors it is also attached to with it and so forth, forming a wide-spanning network that leads to global contraction. We thus propose that the expected number and distribution of crossings between fibers and motors is a determining factor in whether a given randomly arranged collection of actin fibers and molecular motors will contract.

**Figure 2.**
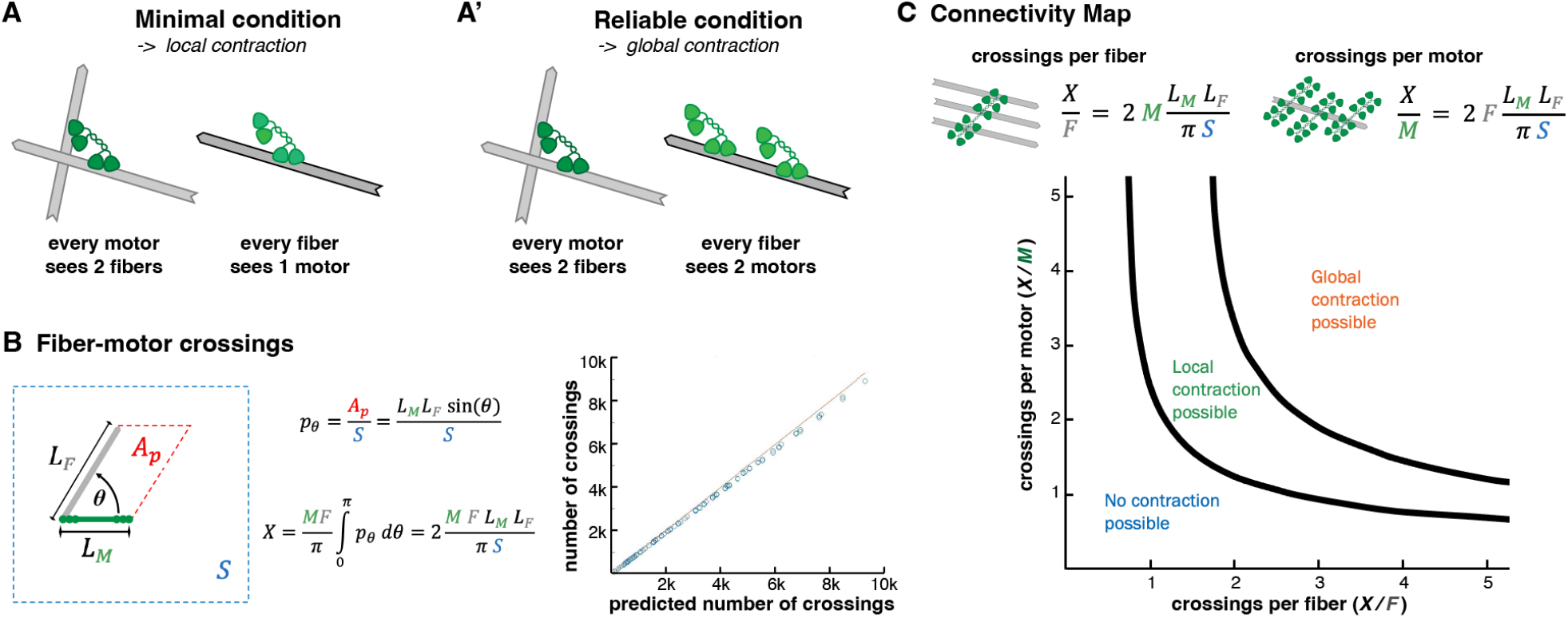
Theory of cytoskeletal connectivity. A) Minimal connectivity condition for the onset of local contraction. A”) Reliable connectivity condition that ensure global contraction is possible. B) Left: If the center of a motor at an angle θ from the fiber is placed inside the parallelogram then the two objects will intersect. Right: Comparison of the measured number of fiber-motor crossings with the number predicted by Eq 3 for a randomly chosen number of fibers and motors. C) Connectivity map as a function of the number of crossings per fiber and crossings per motor with the calculated contours thresholds for the onset of local and global contraction

Assuming a random distribution of *M* motors and *F* fibers on a surface of area *S*, the expected number of fiber-motor crossings (*X*) can be calculated as the product *MF* times the probability *p* that a fiber and a motor of lengths *L*_*F*_ *and L*_*M*_ will intersect. If a motor and a fiber are randomly placed inside this surface, and if we assume that *L*_*F*_ and *L*_*M*_ are much smaller than *S*, the fiber and motor will intersect if and only if the center of the motor lies inside the parallelogram of side lengths *L*_*F*_, *L*_*M*_ and the angle *θ* between both object orientations. Therefore, the probability *p*_*θ*_ of this intersection occurring is just the ratio between this parallelogram area (*A*_*p*_) and *S*:

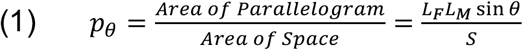

To get the total probability *p* of intersection we can average over all relative angles between the two objects, which are uniformly distributed from *0* to *π*.

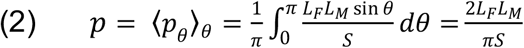

Since each fiber and motor is placed randomly and independently of each other, the total number of expected crossings should be proportional to both the number of fibers *F* and motors *M*.

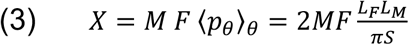

This formula can be confirmed through the computer generation of disordered networks and directly counting the number of geometric intersections. The graph on **Figure 2B** shows the comparison of this actual count and the expected count predicted by Eq 3 for numerous initial configurations with randomly chosen numbers of fibers and motors.

With these calculations we can calculate the density of crossings per each system component as:

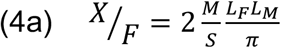

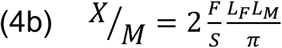

We note that the number of crossings per each component is proportional to the density of the other component and to the lengths of both objects. We also note that while we count geometric crossings, there is not necessarily an active motor head available to physically bind the fiber and motor together. However, given that myosin minifilaments possess many motor heads distributed along its length, we consider that the likelihood of realizing a geometric crossing into a physical one is high.

This approach to counting the number of fiber-motor intersections also serve as the basis to predict the distribution of crossings and the connectivity of the network. For a given fiber, it either intersects or does not each of the *M* motors independently. Therefore, the total number of crossings on that fiber follows the binomial distribution. Eq 2 gives the probability of successful intersection, so it follows that the probability of a fiber having exactly *x* crossings is:

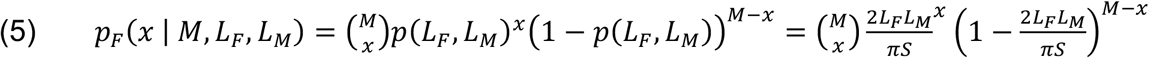

Note that geometrically the role of fibers and motors is symmetric, so these formulas are easily modified to predict motor connections as well.

If the fiber lengths are not uniformly distributed, then we should average over all possible fiber lengths. In living cells, actin fiber lengths are commonly distributed exponentially (Piekenbrock 92). In such a case, the probability of a fiber having *x* intersections with *M* motors is given by:

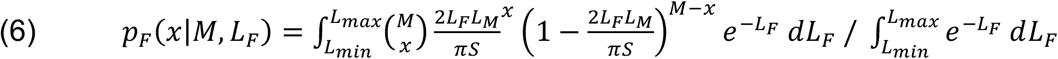

In a true exponential distribution *L*_*min*_ and *L*_*max*_ would be equal to *0* and infinity, respectively, but since in the simulations that we will use to test the theory we can only model a finite number of finite length fibers, we truncate the interval of possible lengths to be between some *L*_*min*_ and *L*_*max*_ values.

The same logic applies to any possible distribution of fiber lengths. Numerical exploration reveals that both versions of the equation depend only on the crossing densities *X/F* and *X/M*, not the individual values independently.

#### Construction of the connectivity map

Next we reintroduce physical behavior to our geometry approach (**Fig 2A,A”**). We can take these distribution probabilities and attempt to predict the potential contractility of a network given an initial count of uniformly distributed fibers and motors. In order for a motor to generate local contraction, said motor must be connected to at least two fibers. The number of motors *M\** that satisfies this condition is:

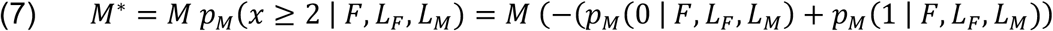

We can then find the probability *P*_*local*_ that a given fiber is connected to at least one of these motors:

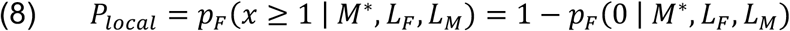

We note that *P*_*local*_ is a function of both *X/F* and *X/M* and forms a connectivity probability map in the *X/F* vs *X/M* space of the likelihood that both conditions shown in **Figure 2A** are met. We hypothesize that a system can only contract locally if a fiber is more likely than not to meet the “*minimal condition*,” i.e., *P*_*local*_ *> 0*.*5*. This defines a curve on the connectivity map that will serve as a threshold that separates the *no contraction* region from the *local contraction* region (**Fig 2C**).

To find a condition that ensures global connectivity, we can take the same approach used to find *P*_*local*_ and impose the “*reliable condition*” (**Fig 2A”**), where a fiber must be connected to at least two motors that in turn are connected to at least two fibers.

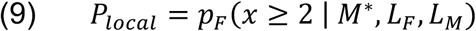

Using the same argument as before, we predict that a system will contract globally if this probability is more likely than not (*P*_*global*_ *> 0*.*5*). This defines a curve on the parameter map that will serve as a connectivity threshold that separates the *local contraction* region from the *global contraction* region (**Fig 2C**).

#### Generalization for different length distributions

The derivation above assumes all fibers have the same length. However, we can broaden our approach to account for any fiber length distribution. Since our probabilities rely on fiber and motor count, which are discrete, we first need to approximate continuous distributions with a finite set of lengths *{L*_*j*_*}* with a corresponding set of probabilities *{q(L*_*j*_*)}*. The number of fibers of each length is thus rounded down so they can be used in a binomial coefficient.

As before we need to find the number of motors attached to at least two fibers. We will make a simplifying assumption here that this can be calculated using the average fiber length ⟨*L*_*F*_⟩:

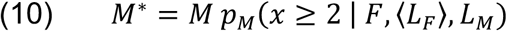

Then for each set of fibers we predict the number of fibers of that length connected to at least one of these motors. We then take a weighted sum over each length and find *P*_*local*_ and *P*_*global*_ as before. As we will show in the results section, a weight of *(L*_*j*_*/⟨L*_*F*_*⟩)*^*2*^ is appropriate. This captures the fact that longer fibers are more likely to cross many motors, thus contributing more to network connectivity. Eq 11 and Eq 12 give contraction probabilities for any fiber length distribution:

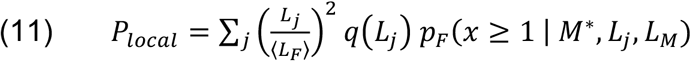

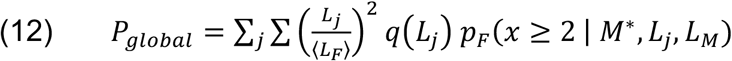

Note that we assumed that motors lengths are uniform, which is a reasonable simplifying assumption as there is little to no variation in motor sizes for any given motor type.

## SIMULATION METHODS AND METRICS

### Computational model

Given the probabilistic nature of our theory, we need to be able to generate a large number of samples with the same input parameters and vary those parameters over a large range of values. This would be difficult and time-consuming to do using experimental setups, so instead we utilize computer simulations. This gives us the ability to finely control the number of each component in the system.

We used the open source cytoskeletal simulation software *CytoSim* (Nedelec 07), which is capable of simulating the thermal environment of the cell as well as the forces generated by the interaction of biological filaments and binding proteins. We identified 3 components essential to understanding polarity sorting contraction: Fibers, myosin motors and crosslinkers (**Fig 3A**). Although the theory derived in this work is generic and applicable to both actin and microtubule cytoskeleton, we focused on the actin cytoskeleton and chose our reference parameters to reflect this system.

**Figure 3.**
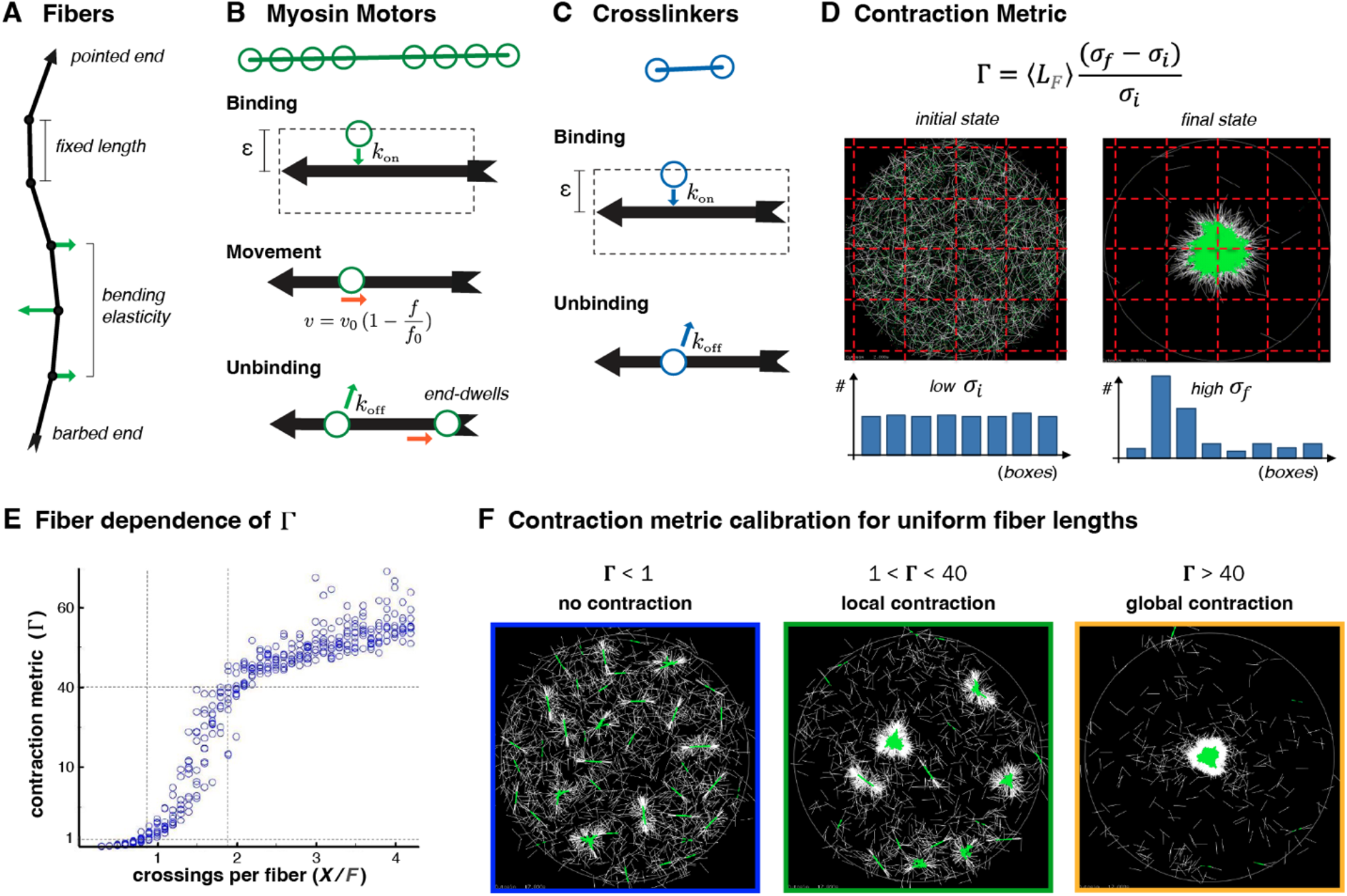
Model and metric. A) A series of rigid segments connected by flexible hinges form a geometrically polarized fiber. B) Myosin minifilaments are modeled as a rod with multiple heads that can bind/unbind fibers and walk along them. C) Crosslinker are modelled as 2 binding heads connected with a spring of resting length of 12 nm. D) Space is divided into boxes and the fold change in spatial standard deviation is used to calculate Γ. E) Γ as a function of crossings per fibers. F) Different ranges of Γ correspond to different contraction outcomes.

Components exist in a flat 2D space mimicking the inner membrane of the cell. The environmental temperature and viscosity match that commonly found in cytoplasm (Daniels 06). The centers of all objects are uniformly, randomly placed inside a circular or periodic space at random orientations with a sufficiently large area size to minimize boundary effects and guarantee that the observed behaviors are not the result of a small space size.

F-actin fibers were modeled as a series of rigid segments connected by flexible hinges, with an adjustable overall rigidity (**Fig 3A**). Fibers are geometrically polarized with a barbed and a pointed end. By default, fiber have a uniform and fixed length of 1 *μm* to mimic a common filament length in an actomyosin network (Burlacu 92) but can be initialized with a more experimentally realistic exponential distribution of lengths.

Molecular motors are designed to model non-muscle myosin II. The backbone of the motor is a 0.8 *μm* long rigid solid, with 12 equally spaced binding heads on each side that cover approximately two-thirds of the backbone”s length (**Fig 3B**). Each head can independently bind to any actin fiber and walk processively towards its barbed end. Upon reaching the barbed end, motors end-dwell there until an unbinding event happens.

Crosslinker were modeled after plastin, which is one of the simplest and most common crosslinkers found *in vivo*. We modeled the crosslinker as a spring with resting length a 0.012 *μm*, with a binding heads at each end that can independently bind to two fibers (Sobral 21, Silva 22) (**Fig 2C**).

A full description of the model with a complete table of parameters is provided in the Supplemental Information.

### Measuring contraction

The most common method of measuring contraction is by looking at the *radius of gyration*. However, this approach has several drawbacks that make it an inappropriate metric here. While it is good at distinguishing between systems that have completely contracted to a single aster and those that have not, it is not able to distinguish between a collection of multiple asters and a stalled contraction. It also requires a complete knowledge of all fiber positions, which is not possible with experimental setups and make the comparison between simulated and experimental results difficult.

To overcome these limitations, we created a new metric based on the percent difference in the spatial standard deviation of fibers. We start by dividing the space into square boxes of equal size. The exact size of each box is not important as long as it is sufficiently larger than the resolution of the object (0.5 *μm* in our case). We then count the number of fiber points positioned in each box, *N*_*kl*_ and calculate *σ* as the spatial standard deviation of {*N*_*kl*_}. As the system contracts and changes from a uniform distribution of fibers to a heterogenous one, *σ* should increase. We define our metric based on the percent change of this difference as:

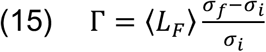

where *σ*_*i*_ and *σ*_*f*_ are the initial and final spatial standard deviation, respectively. The average fiber length <*L*_*F*_> is added to account for the dependency of *σ* on the fiber lengths and allow comparison between simulations where the fiber length distribution is the same, but the average length varies.

## RESULTS

### Metric calibration

Before we can do a systematic exploration of the parameter maps, we must calibrate our contraction metric Γ to identify the correspondence between different metric values and the possible 3 contraction outcomes (**Fig 1B**). When we vary the motor count in our simulations while holding the fiber count steady, we observe an increase in the values of Γ in three distinct phases. Starting at zero (no contraction in the absence of motors), we first observed a slow increase in Γ for low motor counts, followed by a rapid increase at intermediate motor counts, and a sudden decrease in slope for higher motor counts (**Fig 3E**). Inspection of simulation snapshots reveals that the first phase corresponds to the *no contraction* outcome and is characterized as Γ*<1*. The second and third phases correspond to the *local contraction* and the *global contraction* outcomes and are separated at a metric value of Γ*=40*. Careful observations over multiple parameter scans shows that those two metric values (Γ*=1* and *40*) reliably identifies the transition between the 3 contraction outcomes for systems with uniform length distributions (**Fig 3F**). When the length distributions are not uniform, however, the metric must be re-calibrated and we found that for both exponential and gaussian length distribution Γ*=1* still identifies the transition between no contraction to local contraction, but the onset of global contraction is now characterized by Γ*=20* (**Fig S1**).

### Theory correctly predicts contraction outcomes for both rigid and flexible fibers

With a calibrated metric capable of clearly distinguishing contraction outcomes, we explored the full connectivity parameter space by holding the fiber and motor lengths constant and changing their amounts to vary the number of crossings per motor and per fiber. **Figure 4A** shows the full phase space measurements of Γ. The contour lines do not act as perfect dividers but do create guarantees. Any configuration that lands above the *P*_*local*_*=0*.*5* contour has some level of local contraction. Any configuration that lands above the *P*_*global*_*=0*.*5* contour is guaranteed to globally contract. While the theory was derived assuming straight fibers, we found that initializing the fibers with an already relaxed configuration that reflects the persistence length of actin do not alter the results. We attribute this to the fact that the persistence length of F-actin is about an order of magnitude higher than the length values we used (1 *μm*), which is the estimated typical length for actin fibers in the cortex.

**Figure 4.**
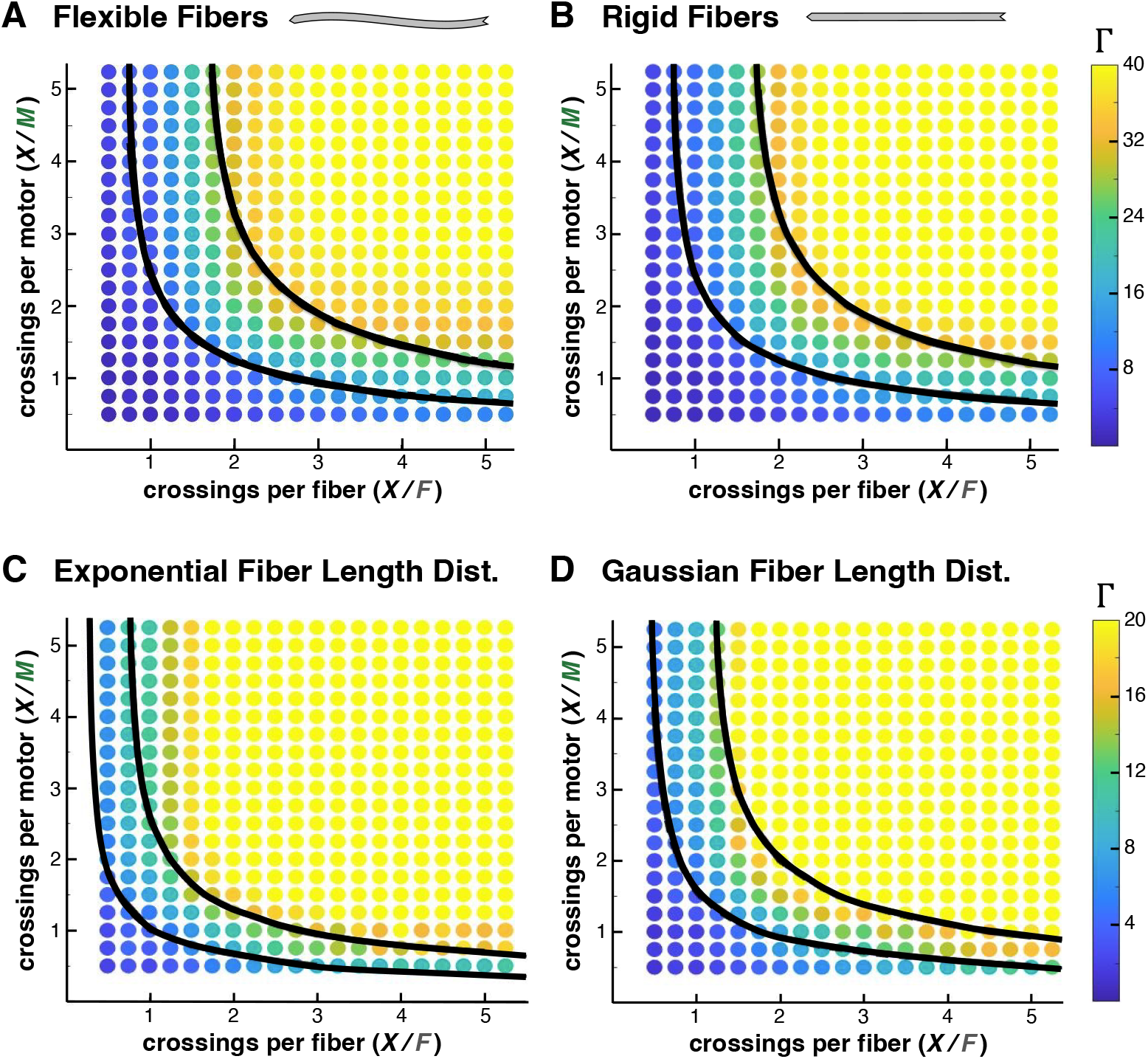
Connectivity parameter maps under different fiber conditions. The contraction metric Γ as a function of crossings per fiber and crossings per motor with (A) uniformly distributed flexible fibers; (B) rigid fibers; and exponential (C) and Gaussian (D) fiber length distributions. Due to the presence of longer fibers that increase crossing likelihoods, the contractile regions in the connectivity maps are expanded for these last two length distributions compared to the uniform case.

We also ran simulations with rigid fibers that are not able to bend. Since polarity sorting is independent of fiber flexibility, we expect that our theory predictions would also hold for this scenario. Our simulation results reveal that the differences in contraction outcomes for flexible and rigid fibers are minimal (**Fig 4A,B**), suggesting that our theory is applicable to both actin and microtubule systems.

### Variations in fiber length distributions

An exponential length distribution for the fibers is a more reasonable model of fibers *in vivo*. Experimental measurements (Piekenbrock 92) have shown actin has this distribution which is the expected result from biopolymers undergoing reversible polymerization (Oosawa 70). Since our theory takes the fiber length distribution as an input, we must first recalculate the *P*_*local*_ and the *P*_*global*_ contour lines using Eqs. 11 and 12, respectively. This leads to similar but shifted separation of the contraction phases when compared to the uniform length distribution (see boundary lines in **Fig 4C**). This right and downward shift towards increased contractility reflects an asymmetry between lengthening and shortening fibers. Long fibers span larger areas and easily accumulate large numbers of crossings leading to more connectivity, and thus fewer components are necessary to drive contraction behaviors. This outweighs the shorter fibers tendency to have fewer crossings, which also contribute less to the contraction of the network and thus do not influence the outcome as much as longer fibers. Our simulations with exponentially distributed fibers confirm our theory predictions of an expanded contractile region and closely align with the predicted boundaries for the onset of local and global contraction (**Fig 4C**). Use of a Gaussian fiber length distribution showed a similar, but less accentuated shift towards increased contractility (**Fig 4D**).

### Results in a periodic space

One difference that does meaningfully change the interpretation and conclusions is attempting contraction in a periodic space. This geometry better reflects conditions where the cytoskeleton network spans over a very large space or the cortex of a cell, where the membrane wraps itself along both directions. Under these geometries high connectivity no longer leads to global contraction. Instead, a well-connected network in a periodic space should not contract at all given that every fiber is connected to its neighbors in all directions and thus feel forces pulling it equally in all directions. We thus expect a different contractility output, where contraction is only possible within the local connectivity region of the parameter space.

Our metric for contraction works the same in a periodic space. Γ*<1* still indicates no contraction, but as **Figure 5A** and **Figure S3** show, this now occurs at both high and low connectivity. Γ*>1* still indicates local contraction leading to the formation of many small but distinct asters. When connectivity is low enough, fibers only interact with nearby neighbors and cannot tell the difference between open versus periodic boundaries. The full phase space matches the understanding that global contraction is no longer possible. While the theory contours still tell us about the probability of local configurations, they no longer translate to global behavior, acting as neither guarantees nor dividers of the space into regions of contractile behavior. Interestingly, we note that most of the local contraction outcome happens along the *P*_*local*_ contour, and diminishes as the crossing per motor increases, but not as much as when the crossing per fiber increases. We attribute this to the higher likelihood that fibers will be left stranded when their amounts are increased relative to the motor counts.

**Figure 5.**
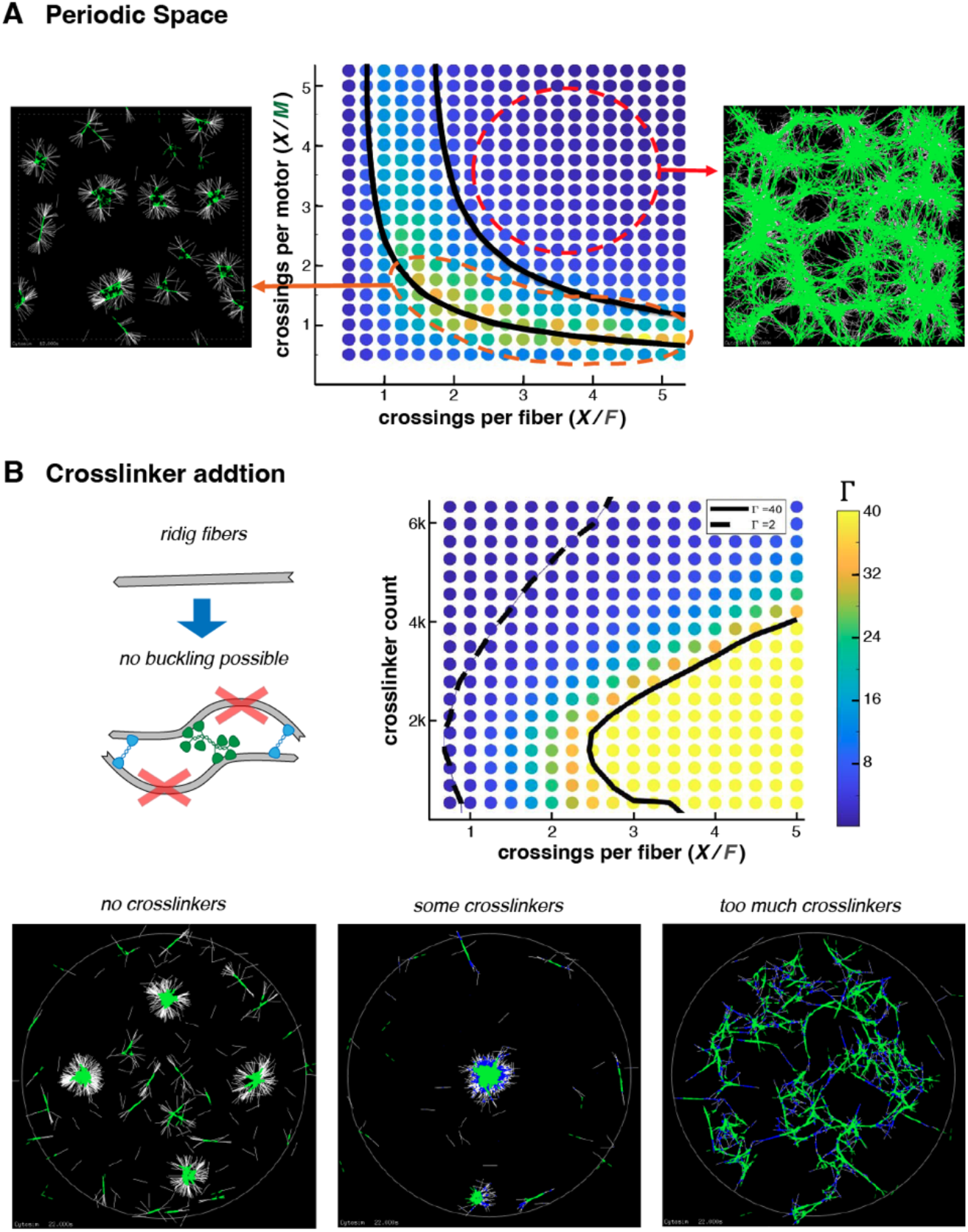
Periodic spaces and crosslinkers. A) Since global contraction in periodic space is no longer possible, the upper contour no longer acts as a meaningful divider of behavior. Instead, local contraction can only be seen along the lower contour. B) A small number of crosslinkers can add enough connectivity to the system to switch the outcome from local to global contraction. Addition of too many crosslinkers eventually lock the network and contraction fails.

### Crosslinkers can rescue global connectivity

Another way of exploring the importance of connectivity is to introduce new types of bonds to the system that, on a local mechanical level, cannot contribute to polarity sorting driven contractions. Crosslinking proteins can bridge pairs of fibers and motors, but do not consume energy or generate forces directly. Their presence also allows for the onset of filament buckling driven contraction (Lenz 12, Murrell 12). In order to study only the connectivity contribution of the crosslinkers, we used rigid fibers that are not able to buckle. This means that the addition of crosslinker connections should harm the contraction of the network by preventing the free reorganization of the fibers by motors.

However, introducing crosslinkers also increases the quantity of fiber-to-fiber connections in the network, and for non-contractile and transition regions, inactive connections may be better than no connections at all.

By introducing crosslinkers, we add a third dimension to our phase space defined by number of crosslinkers *C*. We are not capable of examining every detail of this higher dimensional phase space, so instead we will focus on the absolute number of crosslinkers and select a particular *X/F* vs *C* plane. The graph in **Figure 5B** shows that adding a small number of crosslinkers can rescue systems within the *local connectivity* region to become globally contractile. Fibers that were only bound to one motor and thus not contributing much to the network”s contraction now may be indirectly bound to multiple fibers through a chain of fibers and crosslinkers. While not as reliable, a motor-fiber-crosslinker-fiber-motor chain can drive the same global contraction behavior the motor-fiber-motor chain does. However, as the snapshots in **Figure 5B** show, introducing too many inactive crossings locks the system in its current configuration and stops contraction.

To confirm that connectivity is the necessary explanation for this new contraction, we can compare the system when the fibers are flexible and can contract through the buckling mechanism. Under these conditions the network contraction increases much more dramatically and falls much slower with the addition of crosslinkers when compared to the case where buckling is not possible (**Fig S4**). Since the introduction of a new contraction mechanism creates large changes, the small gains in contraction observed with rigid fibers suggest that low amounts of crosslinkers can enhance contractility by providing higher connectivity to the system.

### Contractility map predicts in vivo network behavior

Next, we check whether the dynamical behavior of the fibers could affect overall network contractility. One way to do this is by adding polymerization and depolymerization of actin fibers in our simulations. Given the complexity of actin fiber dynamics *in vivo*, for which a complete model is still lacking, and many parameters and non-linear effects are still unknown, we decided to first create a simple model for actin dynamics as done in (Sobral 21, Silva 22). We found that if we use reported values of polymerization and depolymerization from *in vitro* experiments (Carlier 17, Shekhar 17) our contraction outcomes still match the theory predictions, but we must use the contour lines for Gaussian fiber distribution as this is the stable fiber distribution under this scenario (**Fig S2**).

The behaviors of actin networks *in vivo*, however, are much more dynamic due to the presence of multiple proteins that nucleate and sever fibers, and also due to non-linear effects that can dramatically increase rates (Arya 24, Reddy 25). Actin fibers are known to undergo fast turnover (Fritzsche 13, Jodoin 15, Goode 23), and recent experiments have shown that myosin minifilaments also disassemble and reassemble *in vivo*. (Kar 23, Chougule 25). One way to model this is by randomly removing and adding fibers and myosin motors to the system at fixed rates, as done in (Belmonte 17). This process tends to disorganize the network while preserving the relative level of connectivity in the system (**Fig 6A**).

**Figure 6.**
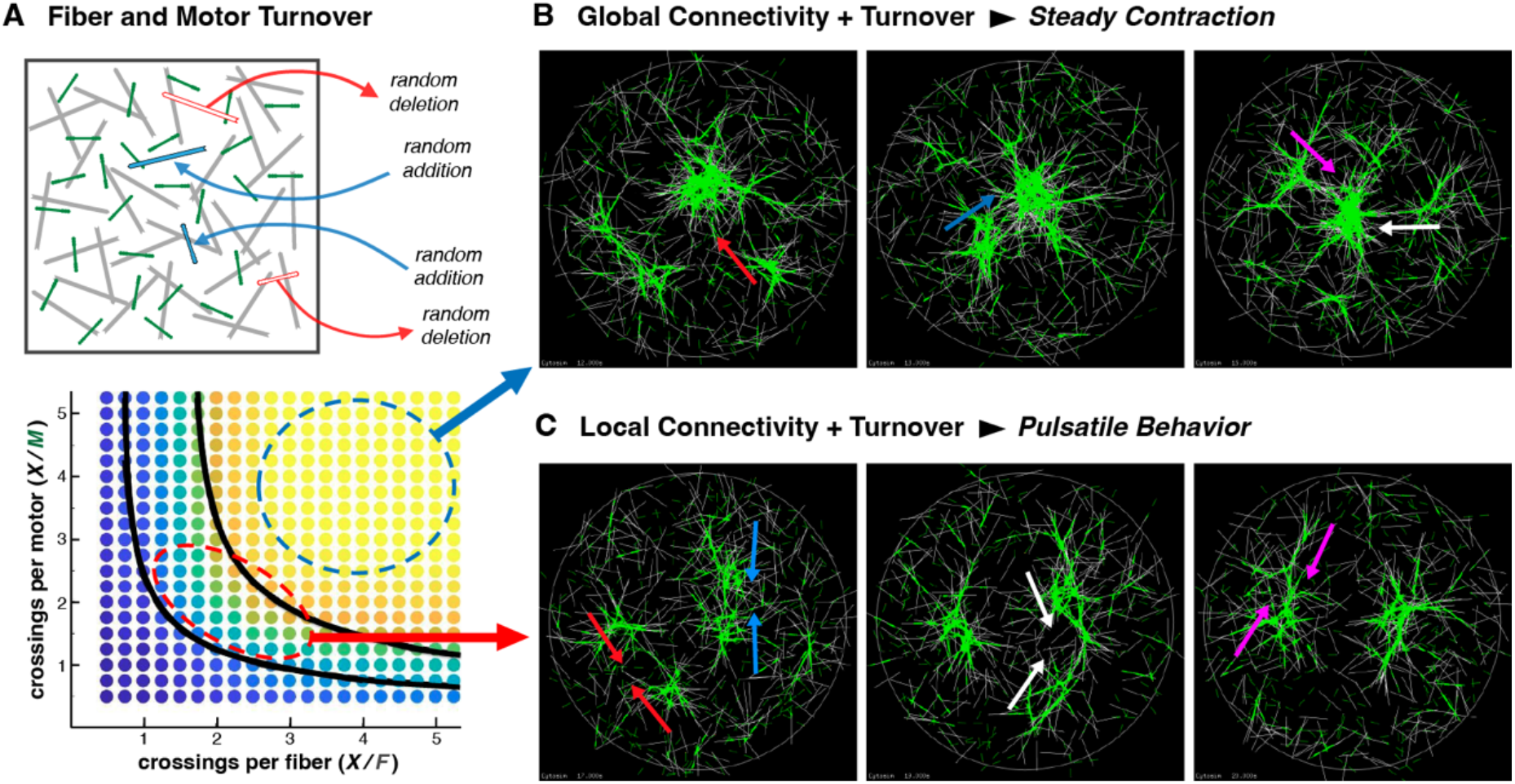
Cytoskeletal turnover and in vivo behavior. A) Fast fiber dynamics is modelled by randomly adding and subtracting different motos and crosslinks at a fixed rate. B) Global connectivity in the presence of turnover leads to a steady contraction of fibers towards the center of the network. Arrows show the direction of accretion of newly formed aster to the center of the network. C) Local connectivity in the presence of turnover leads to pulsatile contraction behavior. Arrows show the merging of asters.

As expected, a poorly connected network remains poorly connected regardless of the turnover rate and no contraction is observed within the low connectivity region of the parameter space. A highly connected network, on the other hand, will stay connected as fibers turnover. An initial large, central aster forms, and as new fibers are added around it, they will form asters that are connected to and attracted towards the central aster through the motor activity (**Fig 5B**). Such steady state contractile influx has been observed in starfish oocytes (Bun et al 18) and in reconstituted actin networks in the presence of turnover (Malik-Garbi 19).

If a system is in the *local connectivity* region, cytoskeletal turnover induces a *pulsatile contraction behavior*. Small asters will form throughout the space, but since the network is only locally connected, the asters will either slowly disappear over time or merge with nearby asters but never form a single contractile center (**Fig 5C**). Over time fibers will be moved from within an aster back to disorganized areas, until the disorganized area is full enough to initiate the formation of new asters. This cytoskeletal behavior has been observed in many biological systems (Sutherland 20), such as during *Drosophila* gastrulation (Martin 09) and *Xenopus* convergent-extension (Kim 11).

## DISCUSSION

The transformation of local interactions of individual fibers and motors into the global behavior of a cytoskeletal network is an area of ongoing interest, with the introduction of ideas from percolation becoming commonplace in investigations. In this work, introduce a novel approach that combines ideas about percolated networks, the role of local motor-fiber interactions and the integration of many fundamental properties of biological fibers and molecular motor proteins. Utilizing tools from probability and geometry, we can predict the number and distribution of force generating connections between fibers and motors based solely on the density of components and their lengths. By examining the local behavior necessary to generate local contraction through the polarity sorting mechanism, we constructed a theory that can predict when a network is sufficiently connected to contract or not, and within the contraction regime, when it will contract locally versus globally.

The connectivity maps that the theory generates depends on fiber length distributions and predicts that wider distributions of fiber lengths favor contraction. We note, however, that those maps are very general and do not need to be recreated for different motor lengths or bigger/smaller systems sizes as both length-scales are already included in the calculation of the connectivity axes, namely crossings per fiber (*X/F=2ML*_*F*_*L*_*M*_*/πS*) and crossings per motor (*X/M=2FL*_*F*_*L*_*M*_*/πS*). Therefore, any cytoskeletal system can be mapped into some location of the connectivity maps provided we know the length of the fibers and motors and their densities.

To test the theory predictions, we developed a new contraction metric Γ based on visual intuition about how asters appear in fiber networks. While ultimately providing a qualitative view contraction, our metric is able to distinguish between the 3 possible network outcomes: no contraction, local contraction and global contraction (**Fig 1B, Fig 3F)**. This metric has many advantages over the commonly used *radius-of-gyration* measurement as it can be used in periodic spaces and for both simulated and experimental images. One disadvantage is that the metric must be calibrated for different fiber length distributions before it can be used to distinguish between the local and global contraction.

Using this new metric and a systematic exploration of the connectivity phase space through computer simulations, we found that the theory predictions were accurate and independent of fiber rigidity, meaning they can be applied to biological networks made up of either actin or microtubules. For microtubule systems molecular motors such as kinesin and dynein are usually bivalent and can only connect two microtubule fibers at the same time, meaning that not all fiber-motos crossing can be converted into an actual connection. However, given that those motors are relatively smaller than the typical myosin minifilaments, we expect that most microtubule motors should cross about 2 fibers, so the theory should still provide accurate predictions for these systems.

We note that while our theory does not account for a bare zone in the myosin motor length, it still matches simulation results with a myosin motor model that has only two thirds of its length covered with motors heads. Attempts at correcting the motor length in our calculations to only cover the active regions yielded wrong predictions. We believe this is because the bare zone lies at the center of the myosin minifilaments, so any fiber crossing at the bare zone will likely find its way to the binding region of the motor as soon as the thermal fluctuations move both objects.

To test whether connectivity was truly fundamental to our approach, we added crosslinkers into a network of rigid fibers which can add connectivity without introducing a new contraction mechanism. We discovered that even for rigid fiber systems, the introduction of a small number of crosslinkers can enhance contractility while a large number will introduce a rigidity percolation effect that stalls contraction. This result highlights the role connectivity plays in the physical behavior of a fiber motor network.

We also tested the importance of connectivity in periodic spaces. The lack of a preferred contraction direction means that a small amount of connectivity is sufficient to drive local contraction, but a well-connected network will try to pull all components in all directions simultaneously and therefore not contract. For these geometries we expect a higher sustained level of tension in the networks at the global connectivity region.

While we have not shown every possible variation and configuration of fiber and motor behaviors, we believe this approach is robust and flexible enough to help predict the behavior of any fiber network that can reasonably be modelled in 2D or recreated in *in vitro* assays. For the last application, our theory can be used as guide to design experiments for which either local or global contraction is the desired result.

The contractility maps may also explain the different modes of contractile behavior seen *in vivo*. We found that in the presence of high actomyosin turnover, simulated systems within the local connectivity region displayed pulsatile contraction, with a continuum formation of multiple asters that either merge with each other or disappear over time. Simulated systems within the global connectivity region, on the other hand, formed a central aster that act as an attractor to which new fibers or newly formed asters adjacent to it are continuously moving. Both outcomes have been observed *in vivo* and our theory predict that the density, or connectivity of actomyosin components must be moderate in cells that display pulsatile behavior, such as the mesodermal cells in *Drosophila* gastrulation or the cells undergoing convergent-extension movements in *Xenopus* embryos (Martin 09, Kim 11). Conversely, it predicts that the density of fibers must be high in the steady contraction of actin filaments after nuclear breakdown in starfish oocytes (Bun 18). Our theory also predicts that pulsatile behavior can happen regardless of boundary conditions, while steady-state contraction must occur in open boundary conditions, where the actin cytoskeleton is not bounded externally as in (Bun 18) and in the *in vitro* experiments with *Xenopus* cytoplasm extracts (Malik-Garbi 19).

Finally, we would like to note that while we explored the connectivity maps by varying only the densities of fibers and motors, we could also have explored the map by changing the length of each object (**Fig 7**). The theory predicts that making a fiber or motor longer while holding the count steady should facilitate contraction by moving the system diagonally towards the global connectivity region. Note that this effect is the same and independent of which object size is being changed, as long as the number of objects is held constant. However, is all fibers are cut into two (as would be if severing proteins are added to the system) then the system will move left in the connectivity map as we combine the effects of shorter fibers (which is down-left movement) and an increase in the fiber count (which is a upward movement), which results in a net loss of connectivity. Therefore, we predict that severing actin fibers may move a globally contractile system (or steady contraction system) into a locally contractile regime (or pulsatile regime) or even to a non-contractile outcome/behavior, depending on where the system lies in the connectivity space.

**Figure 7.**
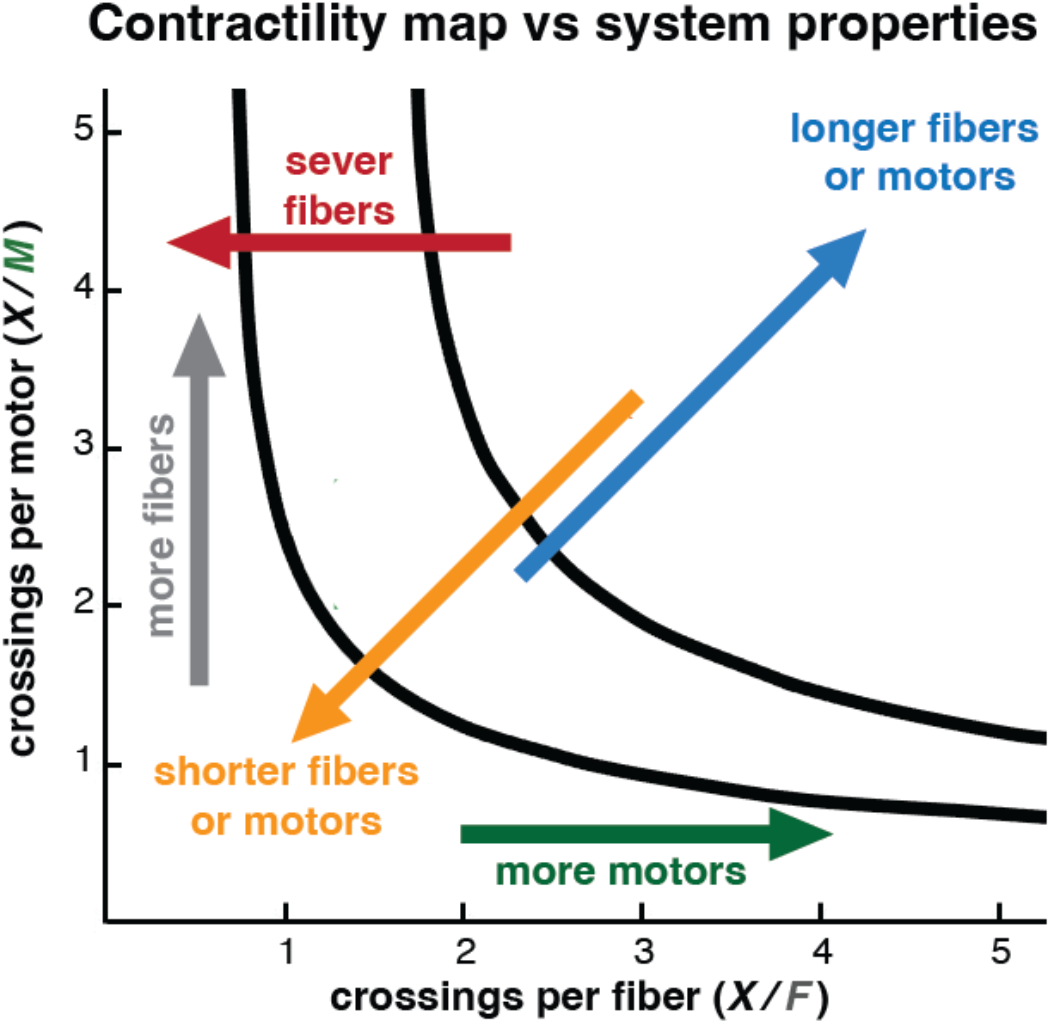
Connectivity map vs system properties. Different changes in system properties leads to different connectivity effects. Addition of more fibers shift the system up the connectivity space (gray arrow), while addition of more motors shift the system right (green arrow). Longer fiber or motors facilitate contraction by shifiting the system diagonally upward-right (blue arrow), while shortening them lead to the reverse effect (orange arrow). Severing fibers combines the effects of fiber shortening and more fibers, resulting in a leftward shif of the system in the connectivity space.

Our theory was developed for the polarity sorting mechanism, which is more tractable than other mechanisms given that it consists of only two elements: fibers and motors. However, we believe that the approach used here can be adapted to the study of connectivity requirements of more complicated contraction mechanisms such as filament buckling, which requires the presence of crosslinkers (or a second type of motor operating at a different speed), and depolymerization end-tracking, which requires a hybrid connector with an end-tracking head and a crosslinker head. These contraction mechanisms would likely add a third dimension to the connectivity maps and the derivation and analysis would be more challenging than the one done here but can potentially bring great insights into the general connectivity requirements of cytoskeletal systems.

## AUTHOR CONTRIBUTIONS

MJN and JMB conceived the work and developed the connectivity theory. MJN run the simulations and analyzed the simulation data under the supervision of JMB. AL and MJN developed and tested the new contraction metric. JMB secured the funding for this work. MJN and JMB wrote the manuscript.

## ACKNOWLEDGEMENTS

We acknowledge the computing resources provided by North Carolina State University High Performance Computing Services Core Facility (RRID:SCR 022168). This work was supported by the North Carolina State University and the National Sciences Foundation (NSF) CAREER award (PHY 2340865).

