## Supplementary Information for "Connectivity and Contraction in Cytoskeletal Networks"

***Supplementary figures***

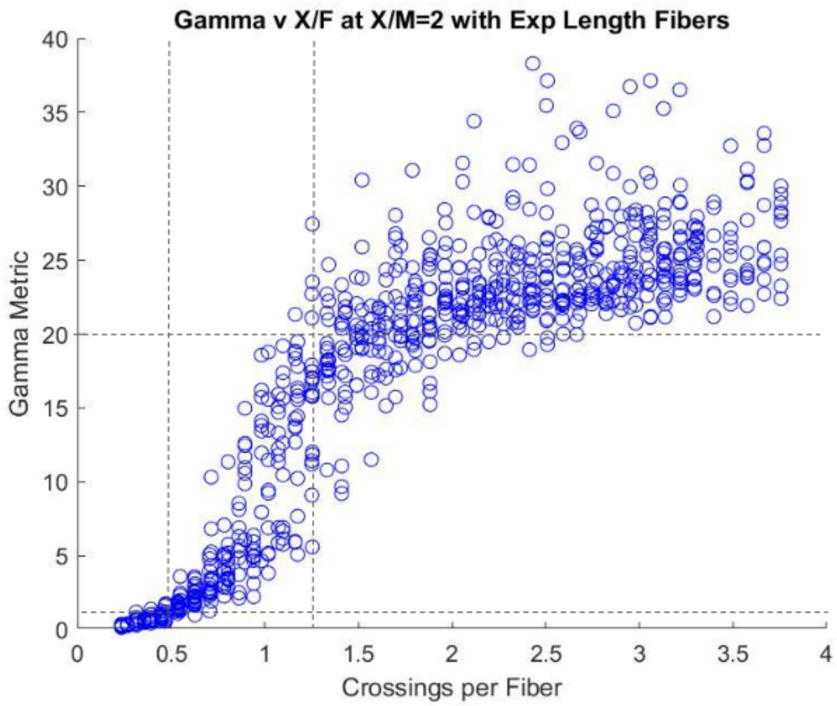


***Figure S1: Γ dependence on X/F for exponential fiber length distribution.*** *The contraction metric Γ increases slowly toward 1 while in the no connectivity region, increases rapidly in the local connectivity region, before changing slope again once entering the global connectivity region. While the variation of the Γ value is higher than for uniform fiber lengths, the basic shape remains. This is unsurprising given there is more input randomness than before. Like in* ***Figure 3E****, the threshold and contour lines intersect near areas of behavioral transition in the Γ curve.*

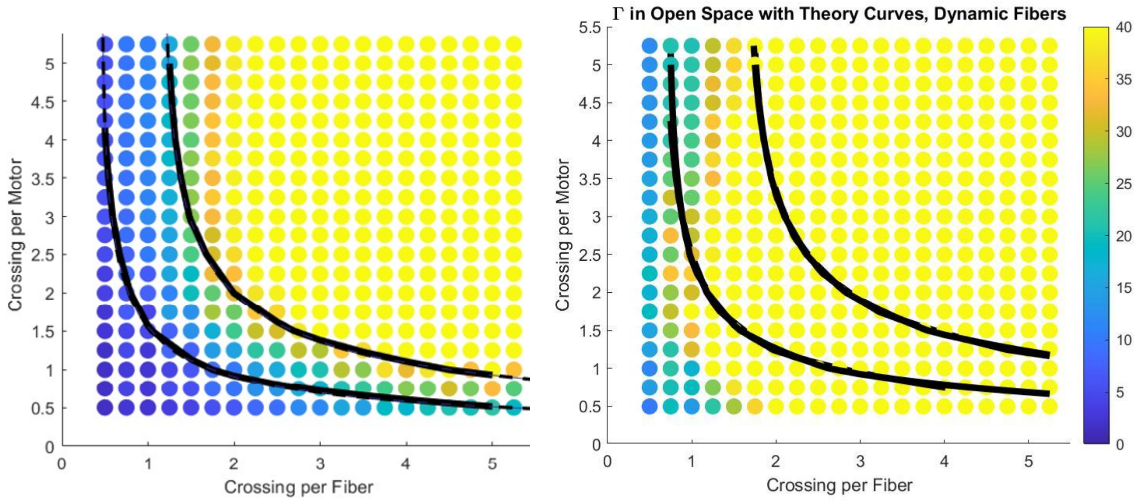


***Figure S2: Dynamic growth connectivity maps.*** *The contraction metric Γ as a function of crossing per fiber X/F and crossing per motor X/M with dynamic fibers. At reasonable growth rates, the overall contractile behavior does not change, and our theory curves still make good guarantors of contraction. However, if the fibers grow unreasonably fast, the theory lines are not predictive anymore.*

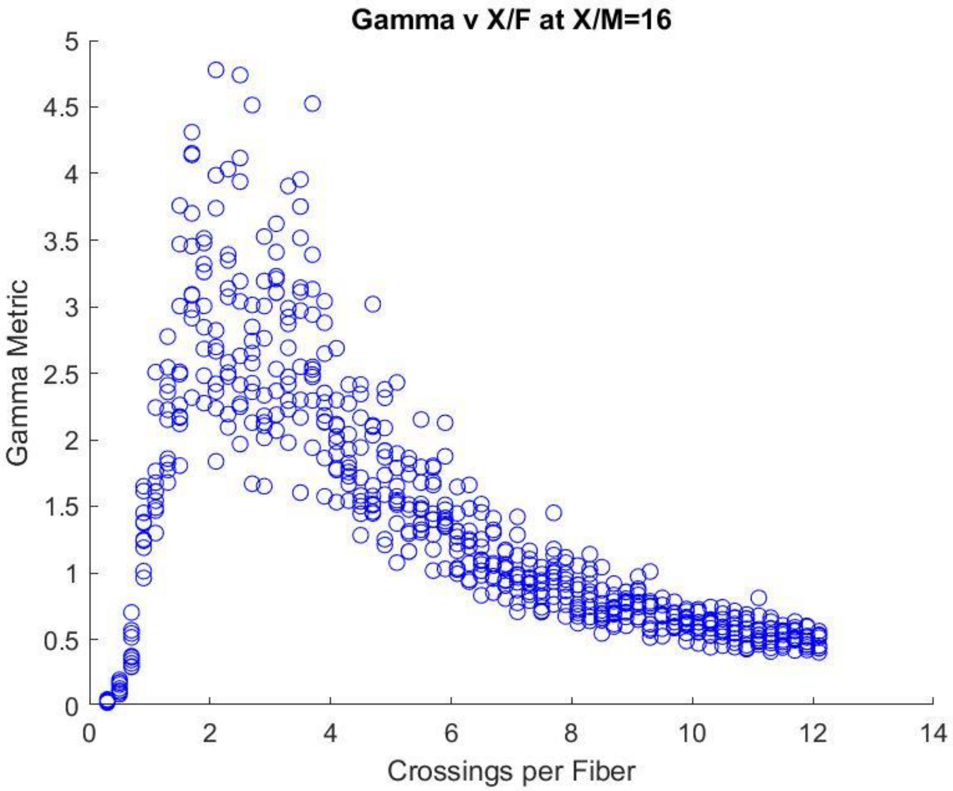


***Figure S3. Γ in a Periodic Space:*** *The contraction metric Γ as a function of crossings per fiber X/F for a fixed density of 16 crossings per motor X/M in a periodic space. Γ increases steadily as it enters the local connectivity region and then decreases as the system becomes connected enough for the boundary conditions to be relevant to the local behavior of fibers. Eventually the metric goes back below 1, indicating the system is so connected it has locked into the initial configuration*.

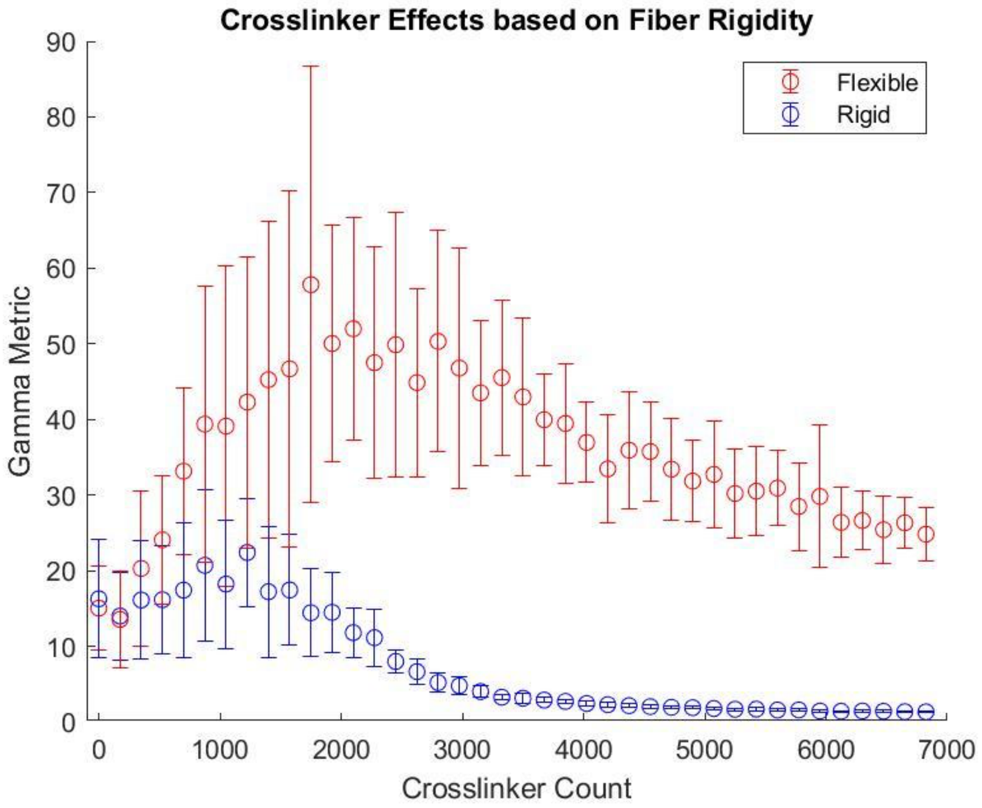


***Figure S4: Γ dependence on addition of crosslinkers.*** *The average with error bars over many trials of contraction metric Γ as a function of crosslinker count a X/M=X/F=1.75 for rigid (blue) and flexible (red) fibers. Crosslinkers help contract both systems through increases in connectivity. Due to the ability to drive contraction through the buckling of flexible fibers, the presence of crosslinkers is far more impactful for flexible fibers.*

### Computational Methods

We simulated the contraction of cytoskeletal networks using the open source engine *CytoSim* (Nedelec 07), which is capable of simulating the thermal environment of the cell as well as the forces generated by the interaction of biological filaments and binding proteins. We identified three components essential to understanding polarity sorting contraction. First is the filaments that define the system. The second is the molecular motors that can independently bind two or more filaments and walk towards a specified filament end. Lastly is the crosslinking proteins, consisting of a body with 2 heads that bind to different filaments. Although the theory derived in this work is generic and applicable to both actin and microtubule cytoskeleton, we focused on the actin cytoskeleton and chose our reference parameters to reflect this system. Components exist in a flat two-dimensional space mimicking the inner membrane of the cell. When possible, component properties are given experimentally accurate values. Otherwise, reasonable ranges are used.

**Filaments:** F-actin were modeled as a series of rigid segments connected by flexible hinges, with an adjustable overall rigidity which we set at 0.075 *pN μm*$^2$ to match experimentally verified values (Gittes 93). Fibers are geometrically polarized with a barbed and a pointed end and by default have a uniform and fixed length of 1 *μm* to mimic a common filament length in an actomyosin network (Burlacu 92), but can be initialized with a more experimentally realistic exponential distribution of lengths. Segments are 0.1 *μm* long in order to allow bending of filaments, but in many simulations we modeled F-actin as a single segment of 1 *μm* to speed up simulation given that filament rigidity was found to not matter in the absence of crosslinkers.

| **Parameter** | **Symbol** | **Value Range (standard value)** |
|---|---|---|
| Rigidity (rigid) | $\mu_{rigid}$ | 22 *pN μm*$^2$ |
| Rigidity (flexible) | $\mu_{flexible}$ | 0.075 *pN μm*$^2$ |
| Segmentation Length | $l_{fiber}$ | 0.1 *μm* |
| Initial Filament Length (initial) | $L_F$ | 0.5-5 *μm* (1 *μm*) |

***Table S1.*** *Parameters for simulating Fibers*

**Dynamic filaments:** Standard filaments do not grow over time. We add dynamics to determine if such effects matter to the polarity sorting contraction mechanism in a similar manner as in (Sobral 21, Silva 22). Dynamic filaments have ends with a growth state and a shrink state where they grow and shrink with pre-defined speeds matching the maximum known growth rates of F-actin *in vitro* (Carlier 17), while keeping the average filaments length roughly constant over time. We also tested extremes by modeling filaments with rates ten times as fast. Each end switches from growth to shrink states at a fixed catastrophe rate and reverts back to a growth state at a fixed rescue rate. We prevented fibers from disappearing by defining a minimum length of 0.03 *μm* where they dwell until a rescue event.

| Parameter | Symbol | Value Range (standard value) |
|---|---|---|
| Minimum Filament Length | $L_{min}$ | .03 *μm* |
| Growing Speed | $v_+$ | 0.08, 0.80 *μm*/s |
| Shrinking Speed | $v_-$ | 0.056, 0.560 *μm*/s |
| Catastrophe Rate | $r_c$ | 4 *Hz* |
| Rescue Rate | $r_r$ | 4 *Hz* |

***Table S2.*** *Parameters for simulating Fibers*

**Non-Muscle Myosin II:** The contraction of fibers into an aster is driven by molecular motors designed to model non-muscle myosin II. The backbone of the motor is a 0.8 *μm* long rigid solid. There are 12 binding heads on each side, equally spaced 0.022 *μm* starting from the ends of the solid towards the center. Therefore, one motor has a total of 24 binding domains covering approximately two-thirds of the backbone's length. This coverage ensures reliable physical connections between the simulated fibers and motors. Each head can bind to any actin fiber within a range of 0.01 *μm* at a rate of 10 *Hz*. Unbinding follows a slip-bound unbinding process (Cortes 20). While bound, heads unbind at a rate $u_m$ of 0.3 *Hz* (Guo 06) modified by the ratio between the total force $f$ exerted on the head and the unbinding force $f_u$, which can be set to infinity to remove the force dependence:

(13) $$u_m = u_{m_0} e^{-\frac{f}{f_u}}$$

While bound to an actin fiber, motor heads walk towards the barbed end of the fiber. The background environment is assumed to have an abundance of the ATP required to fuel walking. The walk is modelled as continuous with a specified maximum speed $v_0$, which is then suppressed by the amount of drag force $f$ on the motor and any other fibers it may be bound:

(14) $$v = v_0 \left(1 - \frac{f}{f_0}\right)$$

Motor movement will cease if total drag reaches a specified stall force value $f_0$, typically around 4 *pN* (Walcott 12). Binding heads follow normal unbinding rules when reaching the end of fibers as opposed to immediately falling off. This end-dwelling property of the motor heads is necessary for polarity sorting mechanism to wok. When attached to a dynamic fiber, if the barbed end of a fiber attempts to grow or shrink while a motor head is attached to the end, the motor head will remain attached.

| **Parameter** | **Symbol** | **Value Range** |
| --- | --- | --- |
| Backbone Length | $L_M$ | 0.8 *μm* |
| Number of Binding Heads | $n$ | 24 |
| Binding Range | $d_{bind}$ | 0.01 *μm* |
| Binding Rate | $b_{motor}$ | 10 *Hz* |
| Unbinding Rate | $u_{motor}$ | 0.03 *Hz* |
| Maximum Motor Speed | $v_0$ | 2 *μm/s* |
| Stall Force | $f_0$ | 4 *pN* |

***Table S3.*** *Parameters for simulating Non-Muscle Myosin II motors*

**Crosslinker:** There is a huge variety of crosslinkers of different lengths and properties (Lieleg 10). For simplicity, our crosslinker were modeled after plastin, which is one of the simplest and most common crosslinkers found *in vivo*. We modeled the crosslinker as a spring of resting length a 0.012 *μm*, with two binding heads at each end and a stiffness of 250 pN/*μm* (Matsudaira 83, Sobral 21, Silva 22). Each head can bind any fiber within 0.01 *μm* at a binding rate of 10 *Hz*. To maximize computational consistency, crosslinkers remain bound once attached to a pair of filaments.

While attached to dynamic filaments, binding heads have a chance to rescue a shrinking fiber. If the fiber attempts to shrink past the location with a bound head, it will be forced into the growth state with a 10% probability, otherwise the binding head will unbind and allow the filament to continue shrinking.

| **Parameter** | **Symbol** | **Value Range** |
| --- | --- | --- |
| Backbone Resting Length | $l_{linker}$ | 0.012 *μm* |
| Spring Stiffness | $k$ | 250 pN / *μm* |
| Binding Range | $d_{bind}$ | 0.01 *μm* |
| Binding Rate | $b_{linker}$ | 10 *Hz* |

| | | |
|---|---|---|
| Unbinding Rate | $u_{linker}$ | 0 *Hz* |
| Rescue Probability | $r_{linker}$ | 0.1 |

***Table S4.*** *Parameters for Modelling Linkers*

**Simulation Space and Initial Conditions:** In order to study polarity sorting in an isolated way and allow our results to be experimentally recreated as simply as possible, we initialized components in a 2D circular space of standard radius 10 *μm*. The environmental temperature and viscosity match that commonly found in cytoplasm (Daniels 06). The centers of all objects are uniformly, randomly placed inside this radius at random orientations. The boundary of the space is open and objects may diffuse outside it. The choice for initial size is a tradeoff between computation time and ensuring contraction effects are not a result of small sample noise or of a too constricted space. This spatial configuration gives a preferred direction for contraction radially inward into one or more asters. To further test our ability to use local conditions to predict global behavior, we also initialize components in a square with periodic boundary conditions. The side length of the square is chosen so that the space has approximately the same area as the circular space. This configuration has no preferred direction of contraction, so we expect that a well connected system would show stalled contraction.

| **Parameter** | **Symbol** | **Value Range** |
|---|---|---|
| Radius | $R_0$ | 10 *μm* |
| Viscosity | $\nu$ | 0.18 $pN\ s / \mu m^2$ |
| Temperature | $kT$ | 0.0042 *pN μm* |
| Fiber Count | *F* | 150-3000 |
| Motor Count | *M* | 150-3000 |
| Crosslinker Count | *C* | 0-30,000 |

***Table S5.*** *Parameters for simulating Space*
